# OPN5 and TRPV4 subserve intrinsic photosensitivity in mouse corneal epithelium

**DOI:** 10.1101/2024.11.14.623473

**Authors:** Luka Lapajne, Monika Lakk, Christopher N. Rudzitis, Shruti Vemaraju, Richard A. Lang, Marko Hawlina, David Križaj

## Abstract

The ocular surface protects the eye from pathogens, mechanical impact and harmful radiation. Excessive exposure of corneal epithelial (CE) layers to ultraviolet B (UVB) photons compromises these protective functions and may be associated with inflammation, pain, vision loss and cancer. We investigated the mechanisms that translate corneal epithelial (CE) transduction of UVB photons into intracellular and inflammatory signaling. Optical imaging in dissociated CECs and intact CE sheets showed rapid, UVB-induced increases in intracellular calcium concentration [Ca^2+^]_i_ that were partially reduced by the TRPV4 antagonist HC067047, removal of extracellular Ca^2+^ and knockdown of the Trpv4 gene, and obliterated by depletion of internal calcium stores. Knockdown of neuropsin and inhibition of phospholipase C signaling markedly reduced the amplitude of the evoked calcium signal. UVB photons promoted release of cytokines and chemokines that included interleukins, metalloproteinases and thymic stromal lymphopoietin (TSLP). CECs are thus intrinsically photosensitive, utilizing a rhabdomeric-like phototransduction pathway cou0led to cytokine release to alert trigeminal afferents and stromal keratocytes about the presence of harmful irradiation and protect the visual system from snow blindness, injury, vision loss and cancer.

## 1. Introduction

Our environment is permeated by ultraviolet (UV) photons that cover ∼2% of the solar spectrum, are invisible to us yet profoundly affect our health (*1–3*). Biological effects of UV radiation are predominantly mediated by the UVB band covering the 280 – 315 nm range. Overexposure may induce skin and eye injuries such as sunburn, snow blindness, *xeroderma pigmentosum*, pain, vision loss, cancers as well as accelerate aging (*4–7*). Due to larger pupils and more transparent ocular media, children are especially vulnerable to UVR with up to 80% of a person’s lifetime exposure to UV light reached before the age of 18 (*8*). Pigmentation (tanning) and sunscreen formulations help prevent UV-induced aging and carcinogenesis (*9–12*) but cannot prevent injuries of the eye.

The visual system is protected from UV radiation by the cornea and the conjunctiva. ∼2% of UVB photons reach the lens and ∼1% the retina, with ∼95% of absorption taking place within the stratified nonkeratinizing corneal epithelium (CE) and anterior stroma (*5, 13, 14*). The CE is particularly vulnerable to UVB injury (*15*), with a single UVB dose sufficient to induce genetic and cellular remodeling (*16–18*) while chronic stimulation forces senescence of the limbal epithelium, increases the risk for *pterygium* and *pingueculae* (conjunctival overgrowth) and climatic droplet keratopathy (corneal opacity) (*19, 20*), and may result in partial or total blindness due to photokeratitis (snow blindness), cataract formation, corneal edema, loss of angiogenic privilege, and ocular melanomas (*4, 20–22*). Photokeratitis, the corneal analog of sunburn, is associated with edema and opacification from damaged CE, stroma and endothelium (*23*). CECs respond to UVB radiation with nuclear translocation of NF-kB (*24*), release of matrix metalloproteinases (MMPs), production of reactive oxygen species, inflammasome activation, mitochondrial dysregulation, CEC sloughing, extracellular matrix (ECM) degradation, barrier disruption, edema formation and afferent overexcitation (*25–31*). UVB-induced CE injury is experienced as cytokine-induced pain mediated by subepithelial afferents from the ophthalmic branch of the trigeminal nerve and may be associated with temporary vision loss (*32, 33*). Despite the pervasiveness and clinical burden associated with UVB -induced pathologies, the identity of UVB sensors within the CE, their downstream effectors and relationship to inflammatory and nociceptive pathways remain poorly understood (*20*).

The past two decades have linked vertebrate light sensing to an astounding complexity of transduction mechanisms in which canonical phototransduction, mediated by OPN1-SW cone opsins and OPN2 (rhodopsin) occurs alongside nonvisual transduction mediated through TRP channels (*34*), cryptochromes (*35*), OPN2 (*36*), OPN3 (panopsin) (*37, 38*), OPN4 (melanopsin) (*34, 39, 40*), and OPN5 (neuropsin) (*41–45*) opsins. TRPV1, TRPA1 and/or TRPV4 channels have been implicated in keratinocyte, melanocyte, fibroblast and skin afferent UV sensing and inflammation (*46–51*) and opsins mediate photosensitivity in keratinocytes, lymphocytes, Jurkat cells, fibroblasts, melanocytes, adipocytes and subsets of retinal, spinal cord and hypothalamic neurons (*37, 41, 52, 53*). Examples of opsin collaborating with TRP channels include blue-light-induced melanogenesis mediated through OPN3 -TRPV1 interactions (*38*) and OPN4 - TRPC6/7 mediated ipRGC phototransduction (*34, 54*). OPN5 contributes to retinal and corneal circadian rhythmicity, wound healing and vascular development (*43, 44, 55*) but it is not known whether OPN5-based signaling coexists with TRP-based photosensing.

Inspired by the report that activation of the polymodal Ca^2+^-permeable channel TRPV4 (Transient Receptor Potential Vanilloid isoform 4) is obligatory for UVB transduction in skin keratinocytes (*47*) and taking into account strong TRPV4 expression in mammalian CEs (*56–58*) and the similarity between molecular and functional properties of keratinocytes and CECs (*59*), we investigated the TRPV4-dependence of UVB sensing in corneal epithelia. We found that CECs are intrinsically photosensitive but mainly rely on rhabdomeric-like signaling to mediate clinically relevant UVB dosing.

## 2. Methods

### 2.1. Ethical approval and animals

Animal handling and experiments followed institutional guidelines set by University of Utah IACUC (Protocol #22-02005) and were conducted in accordance with the National Institutes of Health Guide for the Care and Use of Laboratory Animals, and the ARVO Statement for the Use of Animals in Ophthalmic and Vision Research. C57BL/6, *Trpv4^−/−^* mice (obtained from Dr. Wolfgang Liedtke, Duke University) (*60, 61*) and *Opn5^-/-^* mice (*55*) were maintained in a pathogen-free facility with a 12-hour light/dark cycle and unrestrained access to food and water. KO animals were phenotyped as described previously. Data were gathered from 1- to 3-month-old male and female animals with no noted gender differences.

### 2.2. Reagents

The TRPV4 agonist GSK1016790A (GSK101), antagonist HC-067047 (HC-06) and U-7322 were purchased from Sigma (St. Louis, MO, USA) or Cayman Chemical (Ann Arbor, MI, USA). Cyclopiazonic acid (CPA) was obtained from Tocris (Bristol, UK), and other salts and reagents were obtained from Sigma, VWR (West Chester, PA, USA), Across Organics (Pittsburgh, PA, USA), or ThermoFisher (Waltham, MA, USA).

### 2.3. Tissue preparation

Corneas were dissected from enucleated eyeballs and placed in Dulbecco’s Modification of Eagle’s Medium (DMEM)/F12 (1:1 mixture, GIBCO (Grand Island, NY, USA)/Thermo Fisher) containing Dispase II (15 mg/mL, Sigma) and 1% penicillin/streptomycin for 1 hour at 4°C and an additional hour at room temperature (*56*). Epithelial sheets were peeled off and used *in situ*. Dissociated mouse CECs (mCECs) were incubated in DMEM (GIBCO/Thermo Fisher) containing papain (15 U/mL, Worthington (Columbus, OH, USA)) for 30 minutes, rinsed with D-PBS containing 0.5% bovine serum albumin (BSA, Genesee Scientific) and triturated.

### 2.4. Cytokine detection

Cytokine release from mouse CE was tracked with the C-Series Mouse Cytokine Antibody Array C1000 (RayBiotech, Peachtree Corners, GA, USA), with the membranes, antibody cocktails, buffers and streptavidin used per manufacturer’s instructions. Following dissociation, cells were evenly distributed among vials and DMEM/F12 was added to each vial together with the NPTDase inhibitor ARL 67156 (100 μM) and protease inhibitor cocktail (15 μL/mL). The cells’ exposure to pharmacological agents was 30 min and UVB irradiation 5 min. The membranes were visualized by photographic film and the intensity was quantified using ImageJ software (NIH, Bethesda, MD, USA). The samples were evaluated as duplicates in two independent experiments.

### 2.5. UVB irradiation and optical imaging

Detached corneal epithelial sheets and dissociated cells were placed in the recording chamber (Warner Instruments (Hamden, CT, USA)) and loaded with Fura-2 AM (5-10 µM, Life Technologies) for 30 to 45 minutes. The chamber was placed under an upright Nikon (Tokyo, Japan) microscope with 40x (0.80 NA water) objective. The coverslip at the bottom of the recording chamber was fused quartz as per GE 124 standard (VWR, West Chester, PA, USA) with high 260 – 300 nm transmittance. The UVB light source (UVTOP290H LED, Roithner Lasertechnik, Vienna, Austria) was placed underneath the recording chamber, with irradiation density at the corneal epithelial sheet ∼150 μW/mm^2^. CE sheets were superfused with extracellular saline solution containing (in mM): 133 NaCl, 10 HEPES hemisodium salt, 10 glucose, 2.5 KCl, 2 CaCl2, 1.5 MgCl2, 1.25 NaH2PO4, 1 pyruvic acid, 1 lactic acid, and 0.5 glutathione at pH 7.4 with solution exchanges performed via an electronically controlled multibarrel inlet port (Warner Instruments, Hamden, CT). The UVB wavelength emitted by the LED was 295 nm. For calcium imaging, 340 and 380 nm excitation was delivered via a Xe lamp (Lambda DG-4; Sutter Instruments, Novato, CA, USA) and the emission collected at 510 nm with 14-bit CoolSNAPHQ2 camera (Photometrics, Tucson, AZ, USA) (*56, 62, 63*); image acquisition was paused for the duration of UVB irradiation. Backgrounds were subtracted and F340/F380 ratios computed with NIS-Elements software (Nikon, Lockbourne, OH); ΔR/R (peak F340/F380 – baseline/baseline) was used to quantify the amplitude of Ca^2+^ signals, in which R is the ratio of emission intensity at 510 nm evoked by 340 nm excitation versus emission intensity at 510 nm evoked by 380 nm excitation.

### 2.6. Statistical analysis

Data analysis and statistical tests were performed using Origin Pro 8.5 (Northampton, MA, USA). Unless otherwise stated, data were acquired from at least three independent experiments. Results are given as mean ± SEM. Unpaired sample t-test was used to compare two means, and 1- or 2-way ANOVA with Tukey’s test to analyze three or more means. P > 0.05 = nonsignificant (N.S.), P ≤ 0.05 = *, P ≤ 0.01 = **, P ≤ 0.001 = ***, P ≤ 0.0001 = ****.

## 3. Results

### 3.1. UVB drives CEC release of proinflammatory molecules

UVB -induced inflammation may compromise the integrity of the CE barrier via fibrosis and photokeratitis, and contribute to corneal pain (*18, 30, 57, 64*). To circumvent keratocyte and afferent involvement, we experimentally debrided the CE layer and tracked UVB -dependent cytokine/chemokine release using a chemiluminescence assay. UVB evoked >50% increase in release for 30/96 tested proteins that included interleukins IL-1β (P < 0.001), IL-2β (P < 0.05), IL-17 (P < 0.01), metalloproteinases MMP-3 (P < 0.05), proMMP-9 (P < 0.005), and thymic stromal lymphopoietin (TSLP) (P < 0.001) (Fig. 1A and STable I). Upregulation was observed for the TNF receptor superfamily member 8 (CD30/TNFRS8), Cluster of Differentiation 40 (CD40), Insulin-like Growth Factor-binding Protein 3 (IGF-BP3), leptin, Macrophage Inflammatory Protein (MIP), Monokine Induced by Gamma/chemokine Ligand 9 (MIG/CXCL9), Chemokine (C-C motif) Ligand 5 (CCL5/RANTES), E-selectin, osteopontin, Sonic hedgehog (Shh-N), and vascular endothelial growth factor receptors 1 and 3 (VEGFR1, VEGFR3) proteins (STable I). UVB stimuli thus drive CE-autonomous release of multiple proinflammatory cytokines, chemokines and ECM remodeling enzymes.

**Figure 1.**
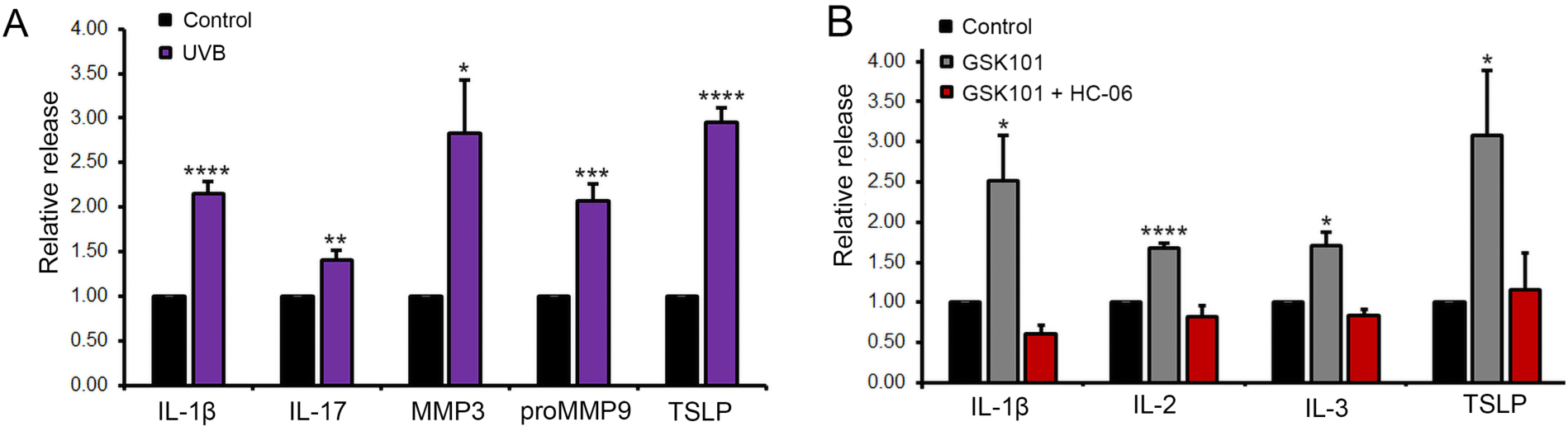
Debrided mouse CE. (A) UVB stimulation promotes release of inflammatory mediators and matrix remodeling enzymes. Debrided mouse CE, chemiluminescent dot assay. Averaged data from 2 independent experiments conducted in duplicate, each sample contained 6 isolated CE. UVB stimulates release of interleukin isoforms, MMPs and TSLP. (B) The TRPV4 agonist GSK101 evokes release of inflammatory mediators and matrix remodeling enzymes. The antagonist HC-06 was co-applied in parallel experiments to validate the specificity of TRPV4 activation. ± SEM, **** P < 0.001, *** P < 0.005, ** P < 0.01, * P < 0.05.

### 3.2. TRPV4 activation induces corneal epithelial cytokine release

Secretion of neuroactive factors from epithelia generally reflects changes in the intracellular concentration of calcium, a 2^nd^ messenger that regulates signaling, proliferation and differentiation, with TRP channels constituting an important venue for Ca^2+^ influx (*65–68*). To test the role of TRPV4 channels in UVB transduction (*47*), debrided mouse CE sheets were exposed to the selective agonist GSK1016790A (GSK101; 25 nM) for 3 min, the duration it takes for the agonist-evoked [Ca^2+^]_i_ response to reach the peak (*56*). GSK101 facilitated >50% release of 24/96 tested proteins (STable 1), with 12 proteins, including IL-1β (P < 0.05), IL-2 (P < 0.0001), IL-3 (P < 0.05) and TSLP (P < 0.05) showing augmented release in response to both GSK101 and UVB stimuli (Fig. 1B) (orange fields; STable 1). Specificity of the TRPV4 mechanism was validated by blocking GSK101-evoked release with the selective antagonist HC067047 (HC-06; 1 μM) (Fig. 1B).

### 3.3. Mouse corneal epithelia are intrinsically photosensitive

To explore the UVB-dependence of CEC Ca^2+^ homeostasis under conditions that minimize contributions from purinergic signaling, afferent, and cell-cell interactions, CEs were dissected from mouse corneas, plated dissociated cells onto UV-permeant quartz glass (*inset* Fig. 2B) and loaded with the ratiometric Ca^2+^ indicator Fura-2-AM (5–10 μM). 3 min exposure to 295 nm light consistently elevated [Ca^2+^]_i_ (Fig. 2Ai-iii), with average peak fluorescence increasing ∼4-fold (from 0.06 ± 0.00 to 0.21 ± 0.01; n = 68 cells, N = 4) (P < 0.0001) (Fig. 2). Hence, mammalian corneal epithelia are intrinsically sensitive to UVB light.

**Figure 2.**
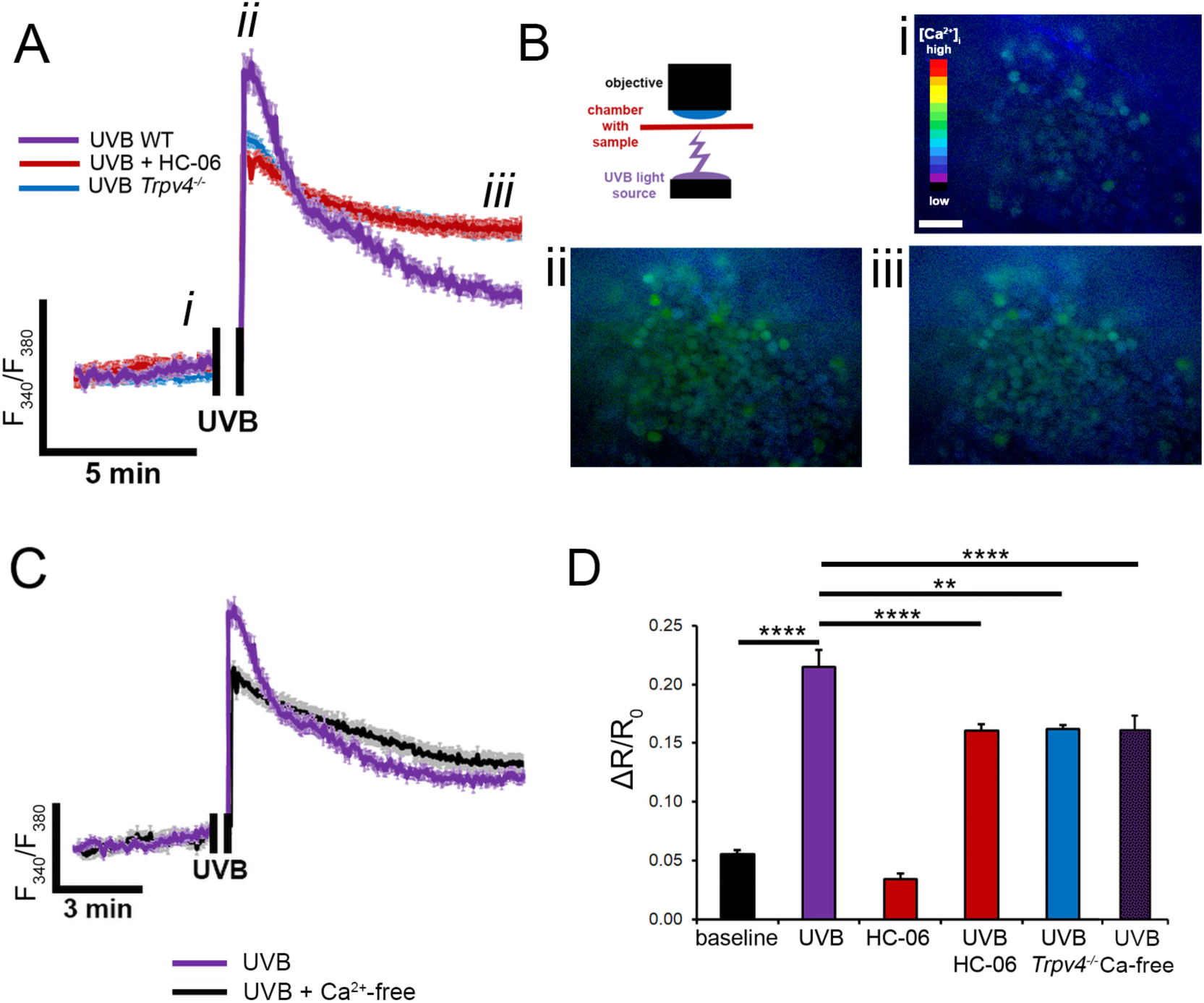
Mouse CE loaded with Fura-2 AM. UVB stimulation induces [Ca^2+^]_i_ increases consisting of peak response that relaxes into a plateau. (A) Representative Ca^2+^ trace from ROIs placed on somata before (i), immediately after (ii) and 10 min after UVB stimulus, with superposed traces from control, HC-06-treated and *Trpv4^-/-^* CE. Fluorescence acquisition was paused during UVB stimulation. (B) Raw images depicting the spatiotemporal changes in the fluorescence ratio at i-iii time-points. Inset, Schematic representation of the experimental configuration. (C) Averaged data show ∼4-fold [Ca^2+^]_i_ increase in UVB-stimulated eyes (P < 0.001), with TRPV4 inhibition (HC-06), deletion (*Trpv4^-/-^*) and Ca^2+^-free saline inducing comparable reductions in [Ca^2+^]_CE_. **** P < 0.001, *** P < 0.005, ** P < 0.01, * P < 0.05.

We tested whether the UVB-dependence of [Ca^2+^]_i_ signals requires TRPV4 involvement using the selective antagonist HC067047 (HC-06; 1 μM), by testing signals in *Trpv4^-/-^* cells and through replacement of control saline with Ca^2+^-free saline. HC-06 reduced the amplitude of the UVB-evoked [Ca^2+^]_i_ response by ∼25% (to 0.16 ± 0.01; N = 4; n = 76; P < 0.0001). CECs isolated from *Trpv4^-/-^* corneas showed comparable reductions in the amplitude of the averaged response (ΔR/R = 0.16 ± 0.01; N = 2; n = 45; P < 0.01) (Fig. 3). The average response in Ca^2+^- free saline was comparable to signals in HC-06-treated and *Trpv4^-/-^* CECs (ΔR/R=0.16 ± 0.01) (N = 2; n = 45, P < 0.01) (Fig. 2A, B), indicating that (i) TRPV4 constitutes the principal component of UVB-induced transmembrane Ca^2+^ influx but (ii) mediates a fraction of the overall calcium response.

**Figure 3.**
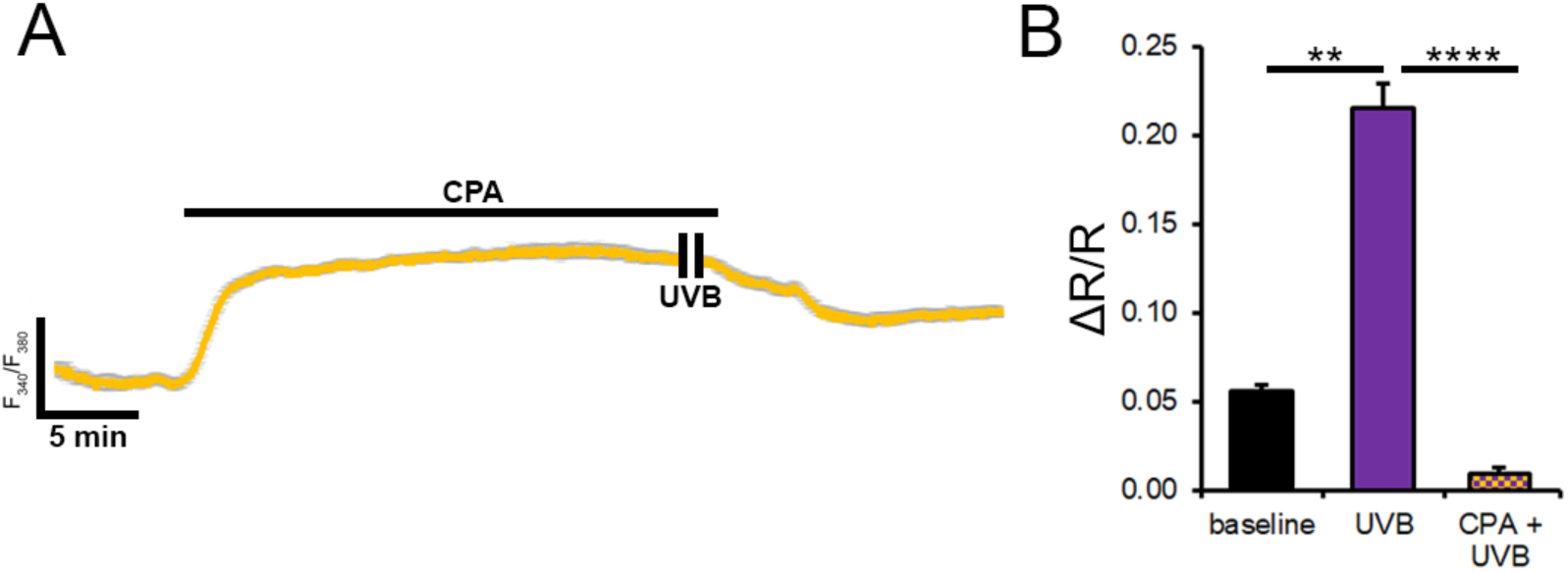
CICR is obligatory for UVB-evoked Ca^2+^ signaling. Dissociated cells. (A) Representative trace of the UVB-evoked Ca^2+^ response in CPA-treated cells. Store depletion is associated with abolition of UVB-evoked. (B) Averaged UVB-evoked response amplitude under control conditions and in the presence of CPA. ** P < 0.01.

### UVB -induced [Ca^2+^]i elevations require release from intracellular stores

The induction of UVB-evoked Ca^2+^ signals in the absence of extracellular Ca^2+^suggests that phototransduction may be upstream from intracellular Ca^2+^ release. We tested the involvement of the ER Ca^2+^ pool using the SERCA transporter blocker cyclopiazonic acid (CPA, 10 μM). As expected (*69, 70*), SERCA inhibition increased cytosolic [Ca^2+^] (to 0.22 ± 0.01; n = 64, P < 0.0001) (Fig. 3, yellow bar). However, depletion of the ER store also abrogated the UVB-evoked response (ΔR/R= 0.01 ± 0.01; n = 64, N = 2; P > 0.05), indicating an obligatory role for Ca^2+^- Induced Ca^2+^ Release (CICR) (Fig. 3).

### 3.4. The CE-intrinsic photoresponse is mediated by neuropsin and phospholipase C signaling

Results depicted in Figures 1-3 show that UVB signaling in the mouse CE requires CICR, with TRPV4 channels mediating a residual fraction of the overall evoked Ca^2+^ signal. We considered involvement of neuropsin, a photopigment encoded by the OPN5 gene that has been implicated in corneal circadian photoentrainment and wound healing (*43, 44*). Comparison of signals evoked by UVB stimuli (20 min) in CECs isolated from wild type and *Opn5^-/-^* animals revealed a significant (P < 0.005) reduction of UVB response amplitude in the KO cohort (Fig. 4). Phospholipase C (PLC) inhibition participates in phototransduction in neurons, melanocytes and keratinocytes (*34, 41, 47, 71*). Consistent with the presence of rhabdomeric signaling in CECs, we found that inhibition of phosphatidylinositol-specific G_q_-coupled phospholipases with U-73122 (1 μM) reduces the amplitude of UVB-evoked by ∼85%, an extent roughly comparable to the effect of OPN5 knockdown (Fig. 4). OPN5 knockdown and PLC inhibition did not affect resting [Ca^2+^]_i_, indicating that OPN5-PLC signaling quiescent in the absence of appropriate photic stimuli.

**Figure 4.**
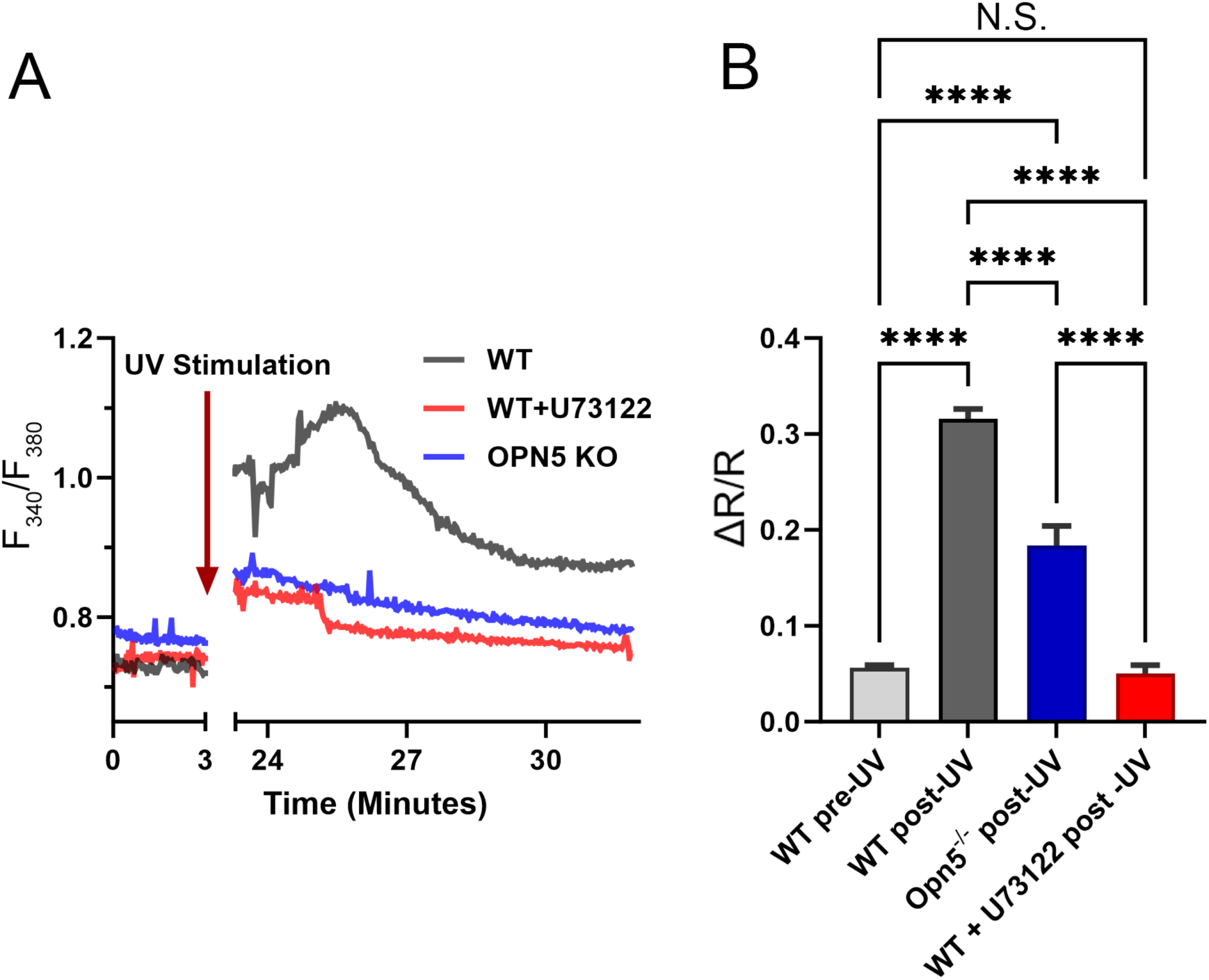
UVB-evoked Ca^2+^ signals require neuropsin and phospholipase C pathways. Dissociated cells. (A) Representative traces for UVB-evoked Ca^2+^ response in control wild type, *Opn5^-/-^* CECs and cells treated with the PLC antagonist U-73122 (1 mM). (B) Averaged data for experiments shown in A. **** P < 0.001

## 4. Discussion

Photokeratitis induced by UVB radiation is characterized by corneal pain and photophobia that protect the eye from photodamage while constraining behavior through temporary vision impairment or loss. The present study identifies epithelial neuropsin and TRPV4 channels as transducers that stimulate release of proinflammatory modulators that may mediate the aversive response to UVB exposure. We show that UVB-induced CE photoresponse requires OPN5 signaling acting in parallel to TRPV4 activation to promote Ca^2+^ signaling and release of proinflammatory cytokines, chemokines and matrix remodeling enzymes (Fig. 5). Acute UVB exposure causes snwo blindness while chronic overexposure can lead to climatic droplet keratopathy, endothelial dysfunction, angiogenesis and oncogenesis resulting from mutagenic cyclobutane pyrimidine dimers and pyrimidine pyrimidone photoproducts, with and propose that the OPN5/TRPV4-PLC-CICR-cytokine axis functions as a rapid alert system that protects vision from high energy photons but could be targeted to mitigate the debilitating effects of inflammation and pain associated with snow blindness and oncogenesis.

**Figure 5.**
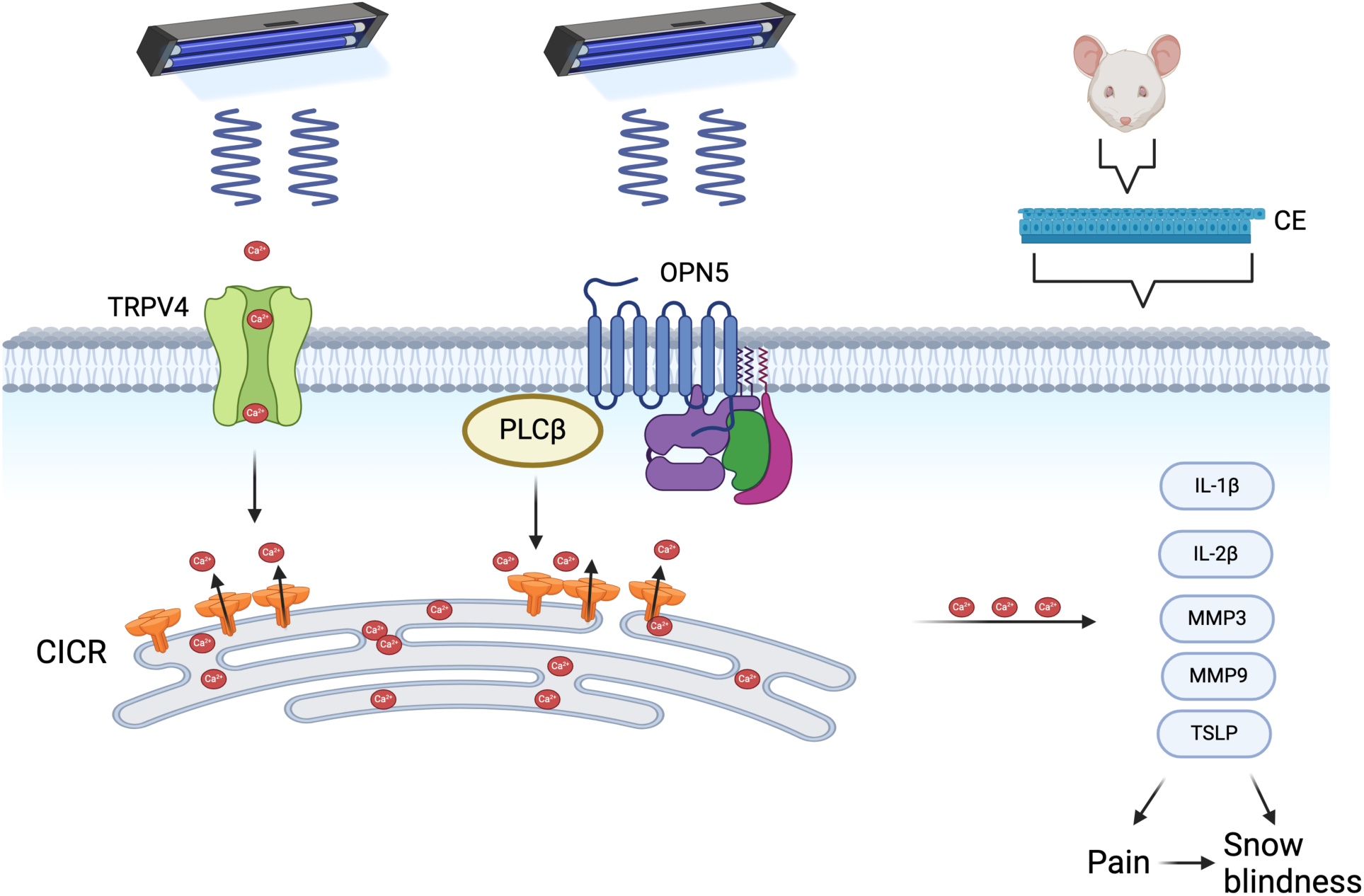
Schema of UVB-induced signaling in the mouse corneal epitheium. Ultraviolet photons stimulate a rhabdomeric-like pathway composed of OPN5, phospholipase -C - G_q_ signaling and attendant CICR mediated through endoplasmic reticulum IP_3_ release channels. [Ca^2+^]_i_ promote gene expression, intracellular signaling and release of inflammatory factors that drive corneal pain and light aversion.

The UVB stimulus utilized in our experiments (150 μW/mm^2^ ; 3 min) corresponds to flux density of ∼27 mJ/mm, approximating ∼20 hrs of outdoor exposure at UV index 10 without sunlight reflection and matching doses used in animal studies (*1, 72–74*) and clinical studies of non-occupational photokeratitis (*29, 75*). Our observation that a single UVB exposure suffices to evoke [Ca^2+^] elevations in in debrided CEs and/or dissociated CECs (i.e., cells separated from stroma and trigeminal feedback) indicates that (i) photosensitivity is an intrinsic property of the corneal epithelium that is (ii) tied to intracellular 2^nd^ messenger signaling. We tested whether CE TRPV4 channels participate in photosensing in a manner that mirrors the TRPV4-dependence of UVB-evoked calcium signaling in mouse keratinocytes (*47*). Consistent with the mechanism proposed by Moore et al. (47), the amplitude of UVB-evoked Ca^2+^ responses was partially reduced by inhibition of TRPV4 channels and deletion of the *Trpv4* gene. Ca^2+^ removal from extracellular saline reduced mCEC responses to an extent comparable to TRPV4 inhibition/knockdown, indicating that (i) TRPV4 constitutes the principal effector of UVB-sensitive transmembrane Ca^2+^ influx, yet (ii) the predominant fraction (∼80%) of the UVB-evoked response is independent of TRPV4. Our observations are consistent with reports that documented the UV-dependence of TRPV1, TRPA1 and TRPV4 activation in multiple types of nonexcitable cell (*48, 51, 76, 77*). The mechanism through which TRP channels respond to UV stimuli is poorly understood, but may involve generation of reactive oxygen species, lipid oxydation, lipid peroxidation, covalent modification of cysteine residues, PLC and/or ET-1 signaling (*46, 47, 50, 78*).

The absorption spectrum of 11-cis-retinal-bound OPN5, a bistable, highly conserved noncanonical opsin (Upton et al., 2022), resembles rod and SWS1/UV (OPN1/2) spectra, with a peak in blue-violet UVA (380 nm *λ*_max_) that extends into the UVB (*λ*_max_ 297 nm) range (*41, 42*). We’ve previously observed that *Opn5^-/-^* mice exhibit normal optokinetic responsiveness (*43*) and SCN-dependent circadian corneal photoentrainment (*44*) whereas debriding induces a CE-intrinsic circadian clock ∼ 3-4 days after corneal wounding (*44*). In contrast to the injury-dependence of CE photoentrainment, our imaging and biochemical experiments suggest that isolated CECs and intact epithelia may be constitutively sensitive to violet light through the OPN5-PLC-CICR axis and auxiliary TRPV4 signaling. Historically considered within the context of bidirectional photoconversion, G_i_ activation and cAMP lowering (*45, 79*), OPN5 photosensing has been recently associated with the G_q_ -PLC - IP_3_ receptor pathway (*80*) that resembles rhabdomeric photosignaling in fly photoreceptors, M1-M7 ipRGCs, melanocytes, choroidal endothelial cells and fibroblasts (*34, 54, 81, 82*). PLC activation cleaves phosphatidylinositol 4,5-bisphosphate (PIP_2_) into IP_3_ and diacylglycerol (DAG) to stimulate calcium release from IP_3_R-sensitive compartments. Consistent with CICR, we found that UVB-evoked [Ca^2+^]_i_ signals largely resisted removal of extracellular Ca^2+^. We confirmed the central role of intracellular release by exposing cells to UVB stimuli during the blockade of SERCA transporters - an intervention that abrogated the evoked Ca^2+^ response – and by stimulating the cells in the in the presence of U73122, which reduced light-evoked response by ∼85% . UVB-evoked calcium signaling in CECs therefore appears to require an intermediary G_q_-PLC step that is linked to CICR and is consistent with the rhabdomeric pathway (Fig. 5). While the precise relationship between OPN5 and TRPV4 transduction its coupling mechanism to store release remain to be delineated in future work, the preservation of the photoresponse under Ca^2+^-free conditions excludes major involvement of Orai/TRPC channels, DAG-sensitive TRPC3/6/7 channels, and Gβγ_q_-adenylate cyclase signaling.

Our findings provide insight into the mechanisms that recruit immune cells to the UVB-injured cornea, stimulate trigeminal afferent fibers and modulate Ca^2+^ -sensitive effector mechanisms such as gene regulation, metabolic regulation, membrane transport, barrier function, and secretion of morphogens and inflammatory mediators. In particular, [Ca^2+^]_i_ elevations and mass release of proinflammatory cytokines and ECM enzymes (e.g., interleukins, MMPs and TSLP) evoked by UVB stimuli and TRPV4 agonis are likely to contribute to the induction and maintenance of corneal pain. With ∼2500 nerve endings per mm^2^, cornea is one of the most densely innervated avascular mammalian tissues. The nociceptive (C) and mechanoreceptive (Aδ) afferents induce aversive responses to mechanical, osmotic, inflammatory and photic stimuli (*57, 83, 84*). The >2-fold increase in the release of IL-1β (Fig. 1) in CEs stimulated with UVB and GSK101 implicate this master regulator of corneal injury in macrophage infiltration, epithelial barrier dysfunction, nociception, angiogenesis, mechanical hyperalgesia and stimulation of downstream MMP, TNF-α, MCP-1 and collagenase pathways (*30, 31, 51, 85*).

Increases in IL-2, IL-17, MMP-3/9, and TSLP release (Fig. 1, Table I) may similarly promote excitation of afferent fibers via monocyte recruitment and ECM degradation (*86*); UVB-evoked release of MMP3/9 stromelysins alone could contribute to stromal thinning, pterygium and corneal neovascularization (*18, 85, 87, 88*). TSLP, an IL-17-like alarmin molecule produced in keratinocytes and epithelia has been linked to immune cell infiltration, innate/adaptive immunity and ocular surface inflammation (*51, 86, 89*). Interestingly, UVB and GSK101 both stimulated release of VEGF molecules, suggesting potential angiogenic functions. Given that keratinocytes also respond to UVB light with secretion of IL-1, IL-6, IL-8, TNFα, TSLP, and MMPs (*51, 88, 90*) it is not inconceivable that noncanonical opsin pathways underlie both snow blindness and sunburn. We propose non-image forming opsins photoentrain peripheral circadian rhythmicity in the absence of SCN inputs (*41*) in order to align tissue responsiveness to daily intensity variation of UV radiation and thus protect the organism from DNA/protein photodamage and cancer (*91*). When these pathways malfunction or are eliminated (as in OPN4/5 KO mice), tissues are at risk for apoptosis, ROS formation, lipid oxydation, inactivated DNA repair, H3K79 methylation and suppression of protective signaling (*71*).

In conclusion, our findings place non-canonical phototransduction in mammalian corneas within the ever expanding range of extraretinal cells and tissues - cranial nerves, heart, skin and GI tract, RGCs, preoptic area of the hypothalamus (OPN4), pineal gland, skin and testis (*36–38, 41, 42, 54, 68, 92, 93*) - that utilize photoreactive opsins to inform and modulate cellular and organismal homeostasis outside of image-forming vision. Constituting a fraction of the overall UV spectrum, UVB radiation is disproportionally phototoxic/genotoxic as the main cause of sunburn, snow blindness and most skin and eye cancers (*3, 8, 94, 95*). Opsin-mediated signaling affords orders-of-magnitude input amplification (*95*) and thus equips cells and organisms with a warning system that alerts them about the need to avoid noxious high-energy photons. Corneal OPN5 presumably works with afferent OPN4 activation (*40, 83*) to drive allodynia, nociception, photophobic behavior and SCN-independent photoentrainment. Additional aspects of corneal OPN5-mediated violet light-sensing might include myopia prevention (*96*). It is worth noting that intraocular pressure experiences circadian rhythmicity that could be modulated by OPN3-5 opsins within the ciliary body (*97*) and that and patients with open angle glaucoma tend to be susceptible to UVA/B damage (*98*). Cornea, ciliary body, ciliary muscle, trabecular meshwork and Schlemm’s canal also express pressure-sensing TRPV4 channels (*56, 62, 63, 99, 100*) and it may be of interest to investigate tissue-intrinsic photosensing and mechanotransduction within the context of chronic ocular hypertension.

## Acknowledgements

We thank Drs. Chia-Yang Liu (Indiana University) for helpful suggestions and Dr. Wolfgang Liedtke (Duke University and Regeneron) for *Trpv4^-/-^* mice.

## AUTHOR CONTRIBUTIONS

LL initiated the study, LL, ML and CR performed the experiments and analyzed the data, SV and RAL provided the OPN5 KO mice, MH and DK obtained funding, LL and DK drafted and wrote the manuscript.

## Supplemental material

**Table 1:**
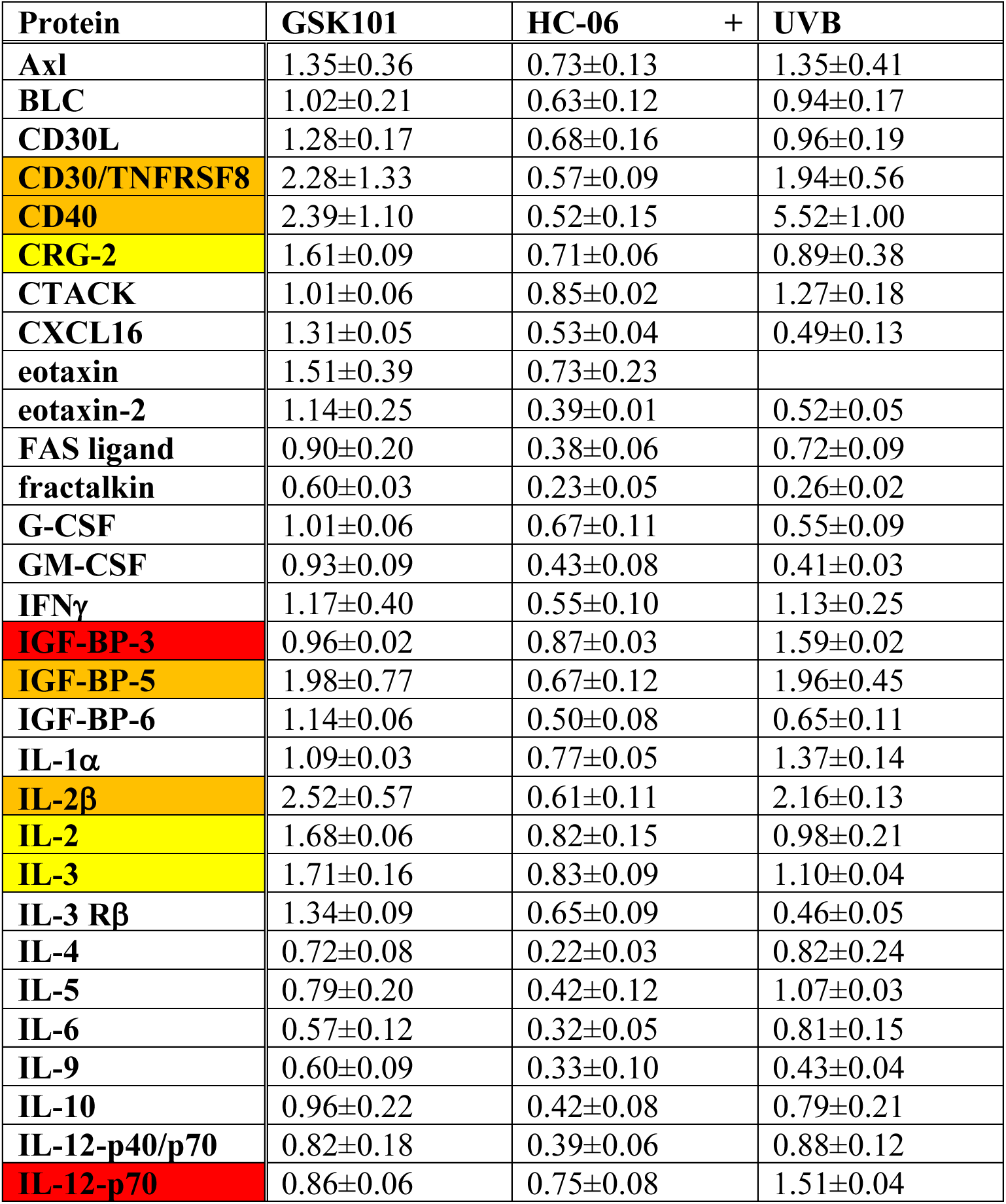
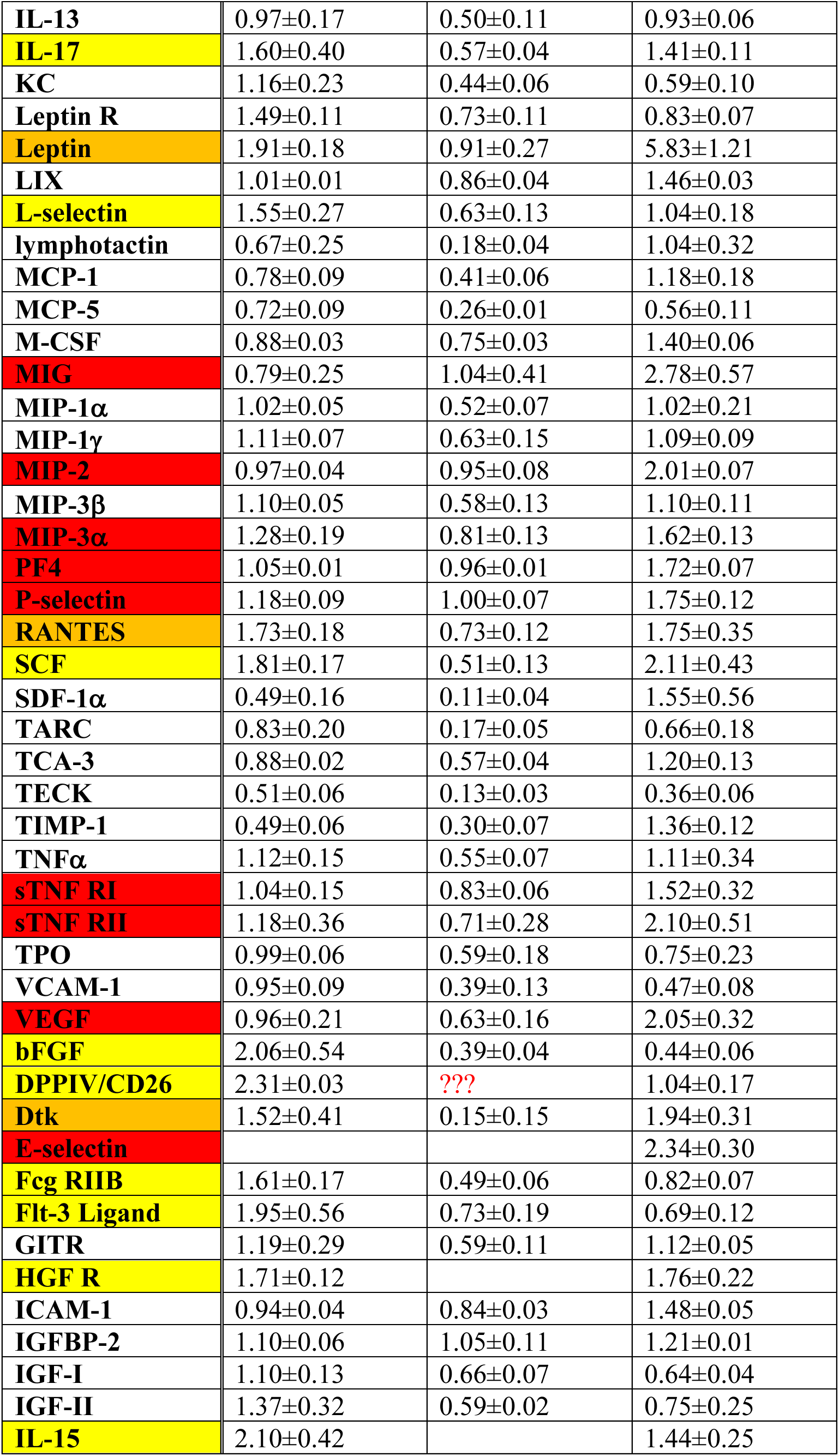
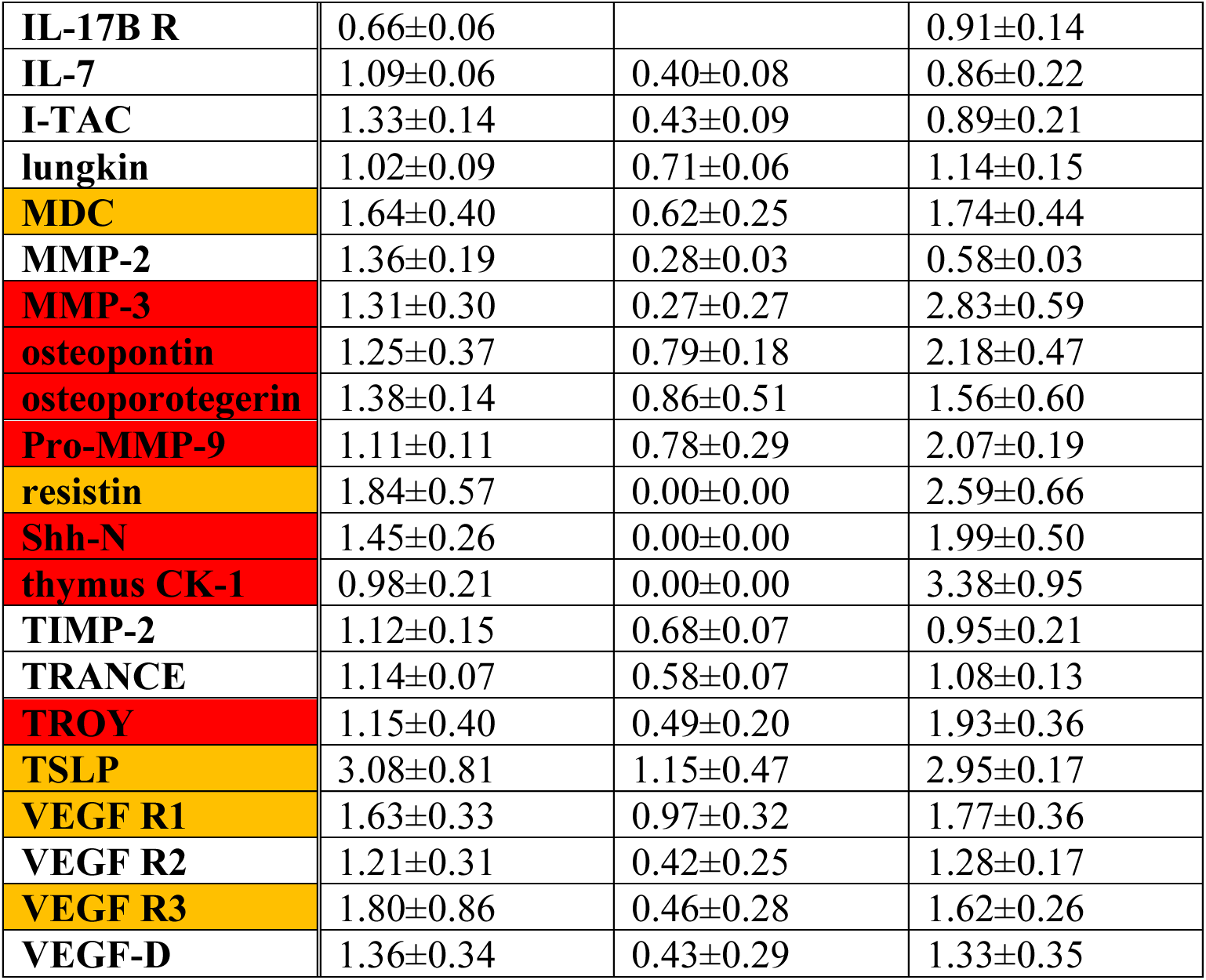
Relative release of 96 proteins following UVB and GSK101 stimulation (control = 1.00). Proteins with >50% increase following 3 min exposure to UVB are marked by red fields. Proteins with >50 release increase following 3 min stimulation with GSK101 (25 nM) labeled with yellow fields; the specificity of the effect was validated with the TRPV4 inhibitor HC-06 (1 μM). Fields with proteins with >50% increase following exposure to UVB and GSK101 are shown in orange. N = 2 independent experiments, n = 2 dot assays/experiment. Average change ± S.E.M.

| Protein | GSK101 | HC-06 + | UVB |
| --- | --- | --- | --- |
| Axl | 1.35±0.36 | 0.73±0.13 | 1.35±0.41 |
| BLC | 1.02±0.21 | 0.63±0.12 | 0.94±0.17 |
| CD30L | 1.28±0.17 | 0.68±0.16 | 0.96±0.19 |
| CD30/TNFRSF8 | 2.28±1.33 | 0.57±0.09 | 1.94±0.56 |
| CD40 | 2.39±1.10 | 0.52±0.15 | 5.52±1.00 |
| CRG-2 | 1.61±0.09 | 0.71±0.06 | 0.89±0.38 |
| CTACK | 1.01±0.06 | 0.85±0.02 | 1.27±0.18 |
| CXCL16 | 1.31±0.05 | 0.53±0.04 | 0.49±0.13 |
| eotaxin | 1.51±0.39 | 0.73±0.23 | |
| eotaxin-2 | 1.14±0.25 | 0.39±0.01 | 0.52±0.05 |
| FAS ligand | 0.90±0.20 | 0.38±0.06 | 0.72±0.09 |
| fractalkin | 0.60±0.03 | 0.23±0.05 | 0.26±0.02 |
| G-CSF | 1.01±0.06 | 0.67±0.11 | 0.55±0.09 |
| GM-CSF | 0.93±0.09 | 0.43±0.08 | 0.41±0.03 |
| IFNγ | 1.17±0.40 | 0.55±0.10 | 1.13±0.25 |
| IGF-BP-3 | 0.96±0.02 | 0.87±0.03 | 1.59±0.02 |
| IGF-BP-5 | 1.98±0.77 | 0.67±0.12 | 1.96±0.45 |
| IGF-BP-6 | 1.14±0.06 | 0.50±0.08 | 0.65±0.11 |
| IL-1α | 1.09±0.03 | 0.77±0.05 | 1.37±0.14 |
| IL-2β | 2.52±0.57 | 0.61±0.11 | 2.16±0.13 |
| IL-2 | 1.68±0.06 | 0.82±0.15 | 0.98±0.21 |
| IL-3 | 1.71±0.16 | 0.83±0.09 | 1.10±0.04 |
| IL-3 Rβ | 1.34±0.09 | 0.65±0.09 | 0.46±0.05 |
| IL-4 | 0.72±0.08 | 0.22±0.03 | 0.82±0.24 |
| IL-5 | 0.79±0.20 | 0.42±0.12 | 1.07±0.03 |
| IL-6 | 0.57±0.12 | 0.32±0.05 | 0.81±0.15 |
| IL-9 | 0.60±0.09 | 0.33±0.10 | 0.43±0.04 |
| IL-10 | 0.96±0.22 | 0.42±0.08 | 0.79±0.21 |
| IL-12-p40/p70 | 0.82±0.18 | 0.39±0.06 | 0.88±0.12 |
| IL-12-p70 | 0.86±0.06 | 0.75±0.08 | 1.51±0.04 |
| **IL-13** | 0.97±0.17 | 0.50±0.11 | 0.93±0.06 |
| **IL-17** | 1.60±0.40 | 0.57±0.04 | 1.41±0.11 |
| **KC** | 1.16±0.23 | 0.44±0.06 | 0.59±0.10 |
| **Leptin R** | 1.49±0.11 | 0.73±0.11 | 0.83±0.07 |
| **Leptin** | 1.91±0.18 | 0.91±0.27 | 5.83±1.21 |
| **LIX** | 1.01±0.01 | 0.86±0.04 | 1.46±0.03 |
| **L-selectin** | 1.55±0.27 | 0.63±0.13 | 1.04±0.18 |
| **lymphotactin** | 0.67±0.25 | 0.18±0.04 | 1.04±0.32 |
| **MCP-1** | 0.78±0.09 | 0.41±0.06 | 1.18±0.18 |
| **MCP-5** | 0.72±0.09 | 0.26±0.01 | 0.56±0.11 |
| **M-CSF** | 0.88±0.03 | 0.75±0.03 | 1.40±0.06 |
| **MIG** | 0.79±0.25 | 1.04±0.41 | 2.78±0.57 |
| **MIP-1α** | 1.02±0.05 | 0.52±0.07 | 1.02±0.21 |
| **MIP-1γ** | 1.11±0.07 | 0.63±0.15 | 1.09±0.09 |
| **MIP-2** | 0.97±0.04 | 0.95±0.08 | 2.01±0.07 |
| **MIP-3β** | 1.10±0.05 | 0.58±0.13 | 1.10±0.11 |
| **MIP-3α** | 1.28±0.19 | 0.81±0.13 | 1.62±0.13 |
| **PF4** | 1.05±0.01 | 0.96±0.01 | 1.72±0.07 |
| **P-selectin** | 1.18±0.09 | 1.00±0.07 | 1.75±0.12 |
| **RANTES** | 1.73±0.18 | 0.73±0.12 | 1.75±0.35 |
| **SCF** | 1.81±0.17 | 0.51±0.13 | 2.11±0.43 |
| **SDF-1α** | 0.49±0.16 | 0.11±0.04 | 1.55±0.56 |
| **TARC** | 0.83±0.20 | 0.17±0.05 | 0.66±0.18 |
| **TCA-3** | 0.88±0.02 | 0.57±0.04 | 1.20±0.13 |
| **TECK** | 0.51±0.06 | 0.13±0.03 | 0.36±0.06 |
| **TIMP-1** | 0.49±0.06 | 0.30±0.07 | 1.36±0.12 |
| **TNFα** | 1.12±0.15 | 0.55±0.07 | 1.11±0.34 |
| **sTNF RI** | 1.04±0.15 | 0.83±0.06 | 1.52±0.32 |
| **sTNF RII** | 1.18±0.36 | 0.71±0.28 | 2.10±0.51 |
| **TPO** | 0.99±0.06 | 0.59±0.18 | 0.75±0.23 |
| **VCAM-1** | 0.95±0.09 | 0.39±0.13 | 0.47±0.08 |
| **VEGF** | 0.96±0.21 | 0.63±0.16 | 2.05±0.32 |
| **bFGF** | 2.06±0.54 | 0.39±0.04 | 0.44±0.06 |
| **DPPIV/CD26** | 2.31±0.03 | ??? | 1.04±0.17 |
| **Dtk** | 1.52±0.41 | 0.15±0.15 | 1.94±0.31 |
| **E-selectin** |  |  | 2.34±0.30 |
| **Fcg RIIB** | 1.61±0.17 | 0.49±0.06 | 0.82±0.07 |
| **Flt-3 Ligand** | 1.95±0.56 | 0.73±0.19 | 0.69±0.12 |
| **GITR** | 1.19±0.29 | 0.59±0.11 | 1.12±0.05 |
| **HGF R** | 1.71±0.12 |  | 1.76±0.22 |
| **ICAM-1** | 0.94±0.04 | 0.84±0.03 | 1.48±0.05 |
| **IGFBP-2** | 1.10±0.06 | 1.05±0.11 | 1.21±0.01 |
| **IGF-I** | 1.10±0.13 | 0.66±0.07 | 0.64±0.04 |
| **IGF-II** | 1.37±0.32 | 0.59±0.02 | 0.75±0.25 |
| **IL-15** | 2.10±0.42 |  | 1.44±0.25 |
| **IL-17B R** | 0.66±0.06 |  | 0.91±0.14 |
| **IL-7** | 1.09±0.06 | 0.40±0.08 | 0.86±0.22 |
| **I-TAC** | 1.33±0.14 | 0.43±0.09 | 0.89±0.21 |
| **lungkin** | 1.02±0.09 | 0.71±0.06 | 1.14±0.15 |
| **MDC** | 1.64±0.40 | 0.62±0.25 | 1.74±0.44 |
| **MMP-2** | 1.36±0.19 | 0.28±0.03 | 0.58±0.03 |
| **MMP-3** | 1.31±0.30 | 0.27±0.27 | 2.83±0.59 |
| **osteopontin** | 1.25±0.37 | 0.79±0.18 | 2.18±0.47 |
| **osteoporotegerin** | 1.38±0.14 | 0.86±0.51 | 1.56±0.60 |
| **Pro-MMP-9** | 1.11±0.11 | 0.78±0.29 | 2.07±0.19 |
| **resistin** | 1.84±0.57 | 0.00±0.00 | 2.59±0.66 |
| **Shh-N** | 1.45±0.26 | 0.00±0.00 | 1.99±0.50 |
| **thymus CK-1** | 0.98±0.21 | 0.00±0.00 | 3.38±0.95 |
| **TIMP-2** | 1.12±0.15 | 0.68±0.07 | 0.95±0.21 |
| **TRANCE** | 1.14±0.07 | 0.58±0.07 | 1.08±0.13 |
| **TROY** | 1.15±0.40 | 0.49±0.20 | 1.93±0.36 |
| **TSLP** | 3.08±0.81 | 1.15±0.47 | 2.95±0.17 |
| **VEGF R1** | 1.63±0.33 | 0.97±0.32 | 1.77±0.36 |
| **VEGF R2** | 1.21±0.31 | 0.42±0.25 | 1.28±0.17 |
| **VEGF R3** | 1.80±0.86 | 0.46±0.28 | 1.62±0.26 |
| **VEGF-D** | 1.36±0.34 | 0.43±0.29 | 1.33±0.35 |

